# Alleviation of Al toxicity by Si is associated with the formation of Al-Si complexes in root tissues of sorghum

**DOI:** 10.1101/229625

**Authors:** Peter M. Kopittke, Alessandra Gianoncelli, George Kourousias, Kathryn Green, Brigid A. McKenna

**Author notes:** Correspondence: Peter M. Kopittke.

## Abstract

Silicon is reported to reduce the toxic effects of Al on root elongation but the *in planta* mechanism by which this occurs remains unclear. Using seedlings of soybean (*Glycine max*) and sorghum (*Sorghum bicolor*), we examined the effect of up to 2 mM Si on root elongation rate (RER) in Al-toxic nutrient solutions. Synchrotron-based low energy X-ray fluorescence (LEXRF) was then used for the *in situ* examination of the distribution of Al and Si within cross-sections cut from the apical tissues of sorghum roots. The addition of Si potentially increased RER in Al-toxic solutions, with RER being up to ca. 0.3 mm h^−1^ (14 %) higher for soybean and ca. 0.2 mm h^−1^ (17 %) higher for sorghum relative to solutions without added Si. This improvement in RER could not be attributed to a change in Al-chemistry of the bulk nutrient solution, nor was it due to a change in the concentration of Al within the apical (0-10 mm) root tissues. Using LEXRF to examine sorghum, it was demonstrated that in roots exposed to both Al and Si, much of the Al was co-located with Si in the mucigel and outer apoplast. These observations suggest that Si reduces the toxicity of Al *in planta* through formation of Al-Si complexes in mucigel and outer cellular tissues, thereby decreasing the binding of Al to the cell wall where it is known to inhibit wall loosening as required for cell elongation.

## 1 Introduction

Aluminum (Al) is the most common metallic element in the earth’s soils (Lindsay, 1979), but at neutral and near-neutral pH values, Al-containing minerals are of low solubility and are not toxic. However, an estimated 30-40 % of all arable soils worldwide are acid (von Uexküll and Mutert,1995; Eswaran et al., 1997), with the solubility of Al being elevated in these acid soils. It has been known for ca. 100 years that elevated levels of soluble Al are highly toxic to plant roots (Veitch,1904; Daikuhara, 1914), reducing root elongation within 5 min by binding strongly to the cell wall and inhibiting loosening (Kopittke et al., 2015). Yet, despite being the most studied trace element in the field of phytotoxicity, much remains unknown regarding the toxic effects of Al.

Silicon, the second most abundant element in the earth’s crust, is regarded as a beneficial element for plant growth (Broadley et al., 2012). Indeed, Si has been shown to enhance resistance to pests and to a range of diseases caused by fungi and bacteria. It has been proposed that this increased resistance is due to a physical effect, with Si forming a layer which impedes penetration by the pest (Broadley et al., 2012). Furthermore, Si has been shown to have a beneficial effect in enhancing resistance and tolerance to various forms of abiotic stress, including salinity, drought, metal toxicity, and temperature stress (Haynes, 2017). It has also been reported that Si alleviates Al toxicity in a range of plant species, including maize (*Zea mays*) (Wang et al., 2004), cotton (*Gossypium hirsutum*) (Li et al., 1989), wheat (*Triticum aestivum*) (Cocker et al., 1998b), sorghum (*Sorghum bicolor*) (Galvez et al., 1987; Hodson and Sangster, 1993), and soybean (*Glycine max*) (Baylis et al., 1994). Three potential mechanisms have been suggested whereby Si has been observed to improve growth in Al-containing solutions (Cocker et al., 1998a). Firstly, in some studies conducted in nutrient solutions, Si (added as sodium metasilicate, for example) was added to solutions already containing Al, with this addition of Si resulting in an increase in solution pH and the precipitation of Al-hydroxides (Li et al., 1996). Secondly, it is also possible that the bioavailability of Al can be reduced directly due to the formation of aluminosilicate complexes within the nutrient solution itself (for example, progressive precipitation of aluminosilicate species) (Barcelo et al., 1993). Finally, some studies have shown that the Si can detoxify Al *in planta* independent of any changes in Al chemistry within the bulk nutrient solution (Cocker et al., 1998b). It is this final (*in planta*) mechanism which is of interest in the present study. However, there remains some uncertainty as to how Si alleviates Al toxicity. One potential mechanism by which Si could alleviate Al toxicity is through the precipitation of aluminosilicate complexes within the plant tissues themselves (Hodson and Evans, 1995; Cocker et al., 1998a; Liang et al., 2001; Liang et al., 2015).

The aim of the present study was to investigate the impact of Si on Al toxicity utilizing seedlings of soybean and sorghum. The effect on Si on root elongation rate (RER) was quantified in Al-toxic nutrient solutions, with changes in soluble (reactive) Al determined colorimetrically. We also used synchrotron-based low energy X-ray fluorescence (LEXRF) to provide information on the cellular and subcellular distribution of Al and Si in root tissues. Due to the ubiquitous nature of Si in soils (Wolt, 1994), gaining an understanding of how Si potentially alleviates Al toxicity is important given that an estimated 30-40 % of all arable soils worldwide are acid.

## 2 Materials and methods

### 2.1 General experimental procedures

Seeds of soybean (cv. Bunya) and sorghum (cv. SF Flourish) were rolled in paper towel and suspended vertically in tap water for either 3 d (soybean) or 2 d (sorghum). Perspex strips, each with seven seedlings, were placed on top of glass beakers containing 650 mL of deionized water with 1 mM CaCl_2_ and 5 μM H_3_BO_3_. Throughout this experiment, we used simple nutrient solutions rather than complete nutrient solutions, as for the examination of Al toxicity, the use of simple nutrient solutions reduces the complexity and uncertainty of Al speciation (Kinraide et al., 1985; Kopittke and Blamey, 2016). However, at a minimum, nutrient solutions must contain Ca and B given that root elongation is inhibited rapidly in their absence due to their low phloem mobility (Kinraide et al.,1985; Goldbach et al., 2001). After 18 h growth in these basal solutions, the seedlings in the Perspex strips were photographed using a digital camera to allow for later measurement of root length before being placed on new beakers containing the treatment solution (see below). The seedlings were photographed after growth for a further 3, 6, 12, 24, 36, and 48 h, with root lengths measured digitally using ImageJ v1.45s (available at: http://imagej.nih.gov/ij/). All plant-growth solutions were continuously aerated.

Experiment 1 aimed to provide initial data on the effect of Si on Al toxicity across a range of Al and Si concentrations and pH values. Given that deionized water may contain trace levels of Si as an impurity [for example, see Rogalla and Romheld (2002)], treatments to which Si were not added (i.e. nominally 0 μM Si) are referred to as ‘-Si’ hereafter. For soybean, 32 treatments were investigated, with four Al concentrations (see below), four Si concentrations (-Si, 0.1, 1, and 2 mM), and two pH values (4.5 and 5.0), each with three replicates. For sorghum, only a single pH value (4.5) was examined (see Results section), with 16 treatments thereby consisting of four Al concentrations (below) with four Si concentrations (-Si, 0.1, 1, and 2 mM), each with three replicates. In addition to a Control (0 μM Al), the Al was supplied at concentrations sufficient to reduce RER by ca. 25, 50, and 75 % over 48 h, being 0, 5, 10, and 30 μM Al for soybean and 0, 5, 10, and 25 μM Al for sorghum. These Al concentrations had been determined from preliminary experiments. Concentrations of Si higher than 2 mM were not investigated as they are not representative of typical soil solutions (Menzies and Bell, 1988; Wolt, 1994). To prepare the treatment solutions, sufficient Na_2_O_3_Si.9H_2_O was dissolved in deionized water with 19 M HCl added to decrease to pH 5.4. For example, to prepare 1 L of solution with 1 mM Si required 0.284 g of Na_2_O_3_Si.9H_2_O and ca. 0.34 mL of 19 M HCl. Next, Ca, B, and Al were added using stock solutions of 0.65 M CaCl_2_, 3.25 mM H_3_BO_3_, and 10 mM AlCl3.6H2O. Finally, 0.1 M HCl was used to adjust pH to the desired value (i.e. either 4.5 or 5.0). Solutions were analyzed to determine the Al concentration, with 10 mL samples collected at the time of transfer (i.e. 0 h) and after 48 h before being filtered (0.22 μm, Millipore), acidified with 20 μL of concentrated HCl, refrigerated at 4 °C, and analyzed using inductively coupled plasma optical emission spectroscopy (ICP-OES). Data for RER were analyzed using a twoway analysis of variance (GenStat v18), with comparisons between means made using Fisher’s protected least significant difference (LSD) test.

Experiment 2 aimed to provide more detailed information for some of the treatments identified in Experiment 1 found to alleviate the toxic effects of Al. For soybean, a total of seven treatments were investigated at pH 4.5, with six Al concentrations (0, 5, 10, 15, 20, and 30 μM Al) without added Si (i.e. -Si) plus one Al concentration (30 μM) at 2 mM Si. For sorghum, a total of eight treatments were investigated at pH 4.5, with six Al concentrations (0, 2.5, 5, 10, 15, and 25 μM Al) without added Si (i.e. -Si) plus two Al concentrations (10 and 25 μM) at 2 mM Si. All treatments were replicated five times. For assessment of the concentration of inorganic monomeric Al concentrations after 0, 24, and 48 h, the pyrocatechol violet (PCV) method was used as described by Kerven et al. (1989). Briefly, 3.0 mL of sample was pipetted into a vial, with 0.20 mL of 0.0375 % PCV reagent added, followed by 1.0 mL of hexamine buffer (prepared as described by Kerven et al. (1989)). Absorbance at 585 nm was measured after 60 s (UV-2600, Shimadzu, Japan). We also used PhreeqcI 3.1.7 (Parkhurst, 2014) to examine the potential formation of aqueous AlH_3_SiO_4_^2+^ (Nordstrom and May, 1996; Pokrovski et al., 1996) using values given in Table 1. Data for RER were analyzed using regression analysis with a Weibull-type function (Taylor et al., 1991).

**Table 1.** Thermodynamic constants used to predict the formation of potential formation of soluble AlH3SiO_4_^2+^ using PhreeqcI 3.1.7 (Parkhurst, 2014). Values were taken from Pokrovski et al. (1996) and Nordstrom and May (1996).

|  | *log K* |
| --- | --- |
| $\text{Al}^{3+} + \text{H}_4\text{SiO}_4^0 \rightleftharpoons \text{AlH}_3\text{SiO}_4^{2+} + \text{H}^+$ | -2.38 |
| $\text{Al}^{3+} + \text{H}_2\text{O} \rightleftharpoons \text{AlOH}^{2+} + \text{H}^+$ | -5.00 |
| $\text{Al}^{3+} + 2\text{H}_2\text{O} \rightleftharpoons \text{Al}(\text{OH})_2^+ + 2\text{H}^+$ | -10.1 |
| $\text{Al}^{3+} + 3\text{H}_2\text{O} \rightleftharpoons \text{Al}(\text{OH})_3^0 + 3\text{H}^+$ | -16.8 |
| $\text{Al}^{3+} + 4\text{H}_2\text{O} \rightleftharpoons \text{Al}(\text{OH})_4^- + 4\text{H}^+$ | -22.99 |

### 2.2 Measurement of bulk Al concentrations in apical root tissues

Bulk concentrations of Al in the root apical tissues were determined for both soybean and sorghum in solutions at pH 4.5. For soybean, the experiment consisted of seven treatments, with six of these seven treatments without added Si (i.e. -Si) (0, 5, 10, 15, 20, and 30 μM Al) and one treatment containing 2 mM Si (30 μM Al). For sorghum, the experiment consisted of eight treatments, with six Al concentrations without added Si (i.e. -Si) (0, 2.5, 5, 10, 15, and 25 μM Al) plus two Al concentrations at 2 mM Si (10 and 25 μM). Each treatment consisted of 75 seedlings suspended above 20 L of nutrient solution containing 1 mM CaCl_2_ and 5 μM H_3_BO_3_ at pH 4.5. The seedlings were grown in this basal solution for 24 h before being moved to new 20 L containers with the Al-containing treatments. The PCV method was used to measure inorganic monomeric Al concentrations after 0, 24, and 48 h. After being exposed to the treatment solutions for 48 h, the seedlings were rinsed in 1 mM CaCl_2_ before the apical tissues (0-10 mm) were excised, briefly placed on filter paper to remove excess moisture, weighed to obtain fresh mass, and dried at 65 °C. Tissues were digested using hydrofluoric acid dissolution (Saito et al., 2005) and analyzed for Al using ICP-OES.

### 2.3 *In situ* analyses of the distribution of Al in root apices

An experiment was conducted to investigate the distribution of Al and Si in root apices of sorghum on a cellular and sub-cellular scale. Sorghum seedlings were grown for 18 h in basal solutions (1 mM CaCl_2_ and 5 μM H_3_BO_3_) at pH 4.5 before being transferred to solutions containing 1 mM CaCl_2_, 5 μM H_3_BO_3_, 25 μM Al, and either -Si or 2 mM Si. After growth for 48 h in these Al-containing solutions, roots were briefly rinsed in 1 mM CaCl_2_ (pH 4.5) and a 200-μm transverse section was cut 5 mm from the apex (this corresponding to the elongation zone). The sections were placed in planchettes filled with hexadecane, and frozen in a high pressure freezer (Leica EM PACT2 with a Leica EM RTS). The high pressure freezing was used to ensure rapid freezing, which occurs within milliseconds. The planchettes were split apart and stored under liquid nitrogen before freeze substitution (Leica EM AFS2) in 2% (v/v) glutaraldehyde in acetone at -90°C for 48 h, warming to 20°C, washing in ethanol, infiltration with LR White Resin, and polymerization. After storage at ambient temperature, a Reichert Ultracut Microtome was used to cut 5-μm thick sections, which were placed on 4-μm thick Ultralene Film.

The LEXRF measurements were conducted using the TwinMic beamline (Gianoncelli et al., 2016) at ELETTRA (BL 1.1L) which has eight Si-drift detectors in an annular back-scattering configuration positioned around the specimen (Gianoncelli et al., 2009). In LEXRF mode, the selected regions were scanned with 1.7 keV excitation energy with a 0.6 μm step size (pixel) and a dwell time of 8 s per pixel. Each individual map was 60 × 60 μm (100 × 100 pixels) with scans taking ca. 22 h to complete. Given that the diameter of the root cylinder was ca. 0.5 mm, it was possible to scan only a small proportion of the total cross-sectional area. For each sample, the area selected to be scanned (60 × 60 μm) was the rhizodermis and outer cortex – this being the area in which Al initially accumulates and where Al is known to rapidly exert its toxic effect (Lazof et al., 1996; Marienfeld et al., 2000; Kopittke et al., 2015). The LEXRF spectra were fitted using PyMCA v4.7.3 (Sole et al., 2007). It was because of the long scan duration (ca. 22 h per scan) that only sorghum was examined by LEXRF, with insufficient time to also examine soybean. Furthermore, although it is known that Si can accumulate at the endodermis in roots of sorghum (Sangster and Parry, 1976; Hodson and Sangster, 1993), we only examined the rhizodermis and outer cortex in the present study given that this is where Al accumulates (and we were interested in Al-Si interactions) and we had only limited beamtime.

## 3 Results

### 3.1 Effects of Si on root elongation rate and soluble Al

Experiment 1 aimed to examine the effect of Si on the rhizotoxicity of Al by examining a range of Al and Si concentrations at two pH values. In all three instances, the interaction between Al and Si was not significant (*P* > 0.05), indicating that the pattern of change in RER upon the addition of Si was similar regardless of the Al concentration (Figure 1). Thus, the LSD values presented allow comparison between the different Si concentrations at any given Al concentration. As expected, the extent to which Si alleviated Al toxicity increased with increasing Si concentration. Indeed, at pH 4.5, the addition of 0.1 mM Si typically did not significantly improve growth in Al-toxic solutions (other than for 10 μM Al for sorghum, Figure 1B), whilst 2 mM Si significantly improved growth in all Al-toxic solutions at pH 4.5 by ca. 0.2-0.4 mm h^−1^ (Figure 1A,B). For roots of soybean grown at pH 5.0, it was noted that Al was less toxic than at pH 4.5, even in the absence of Si (Figure 1C). For example, for soybean grown at pH 4.5, the addition of 5 μM Al (-Si) resulted in a decrease in RER from 2.2 to 1.6 mm h^−1^, yet at pH 5.0 RER was 2.1 mm h^−1^ at 0 μM Al and 2.0 mm h^−1^ at 5 μM Al (Figure 1C). The improved root growth at pH 5.0 is presumably due to the precipitation of Al in these supersaturated solutions (see modelling of Kopittke and Blamey (2016)), and so these seemingly unstable solutions at pH 5.0 were not investigated further. Based upon these results from Experiment 1 (Figure 1), three Si-containing treatments were selected for further investigation, being soybean with 30 μM Al plus 2 mM Si, and sorghum with either 10 or 25 μM plus 2 mM Si.

**Figure 1.**
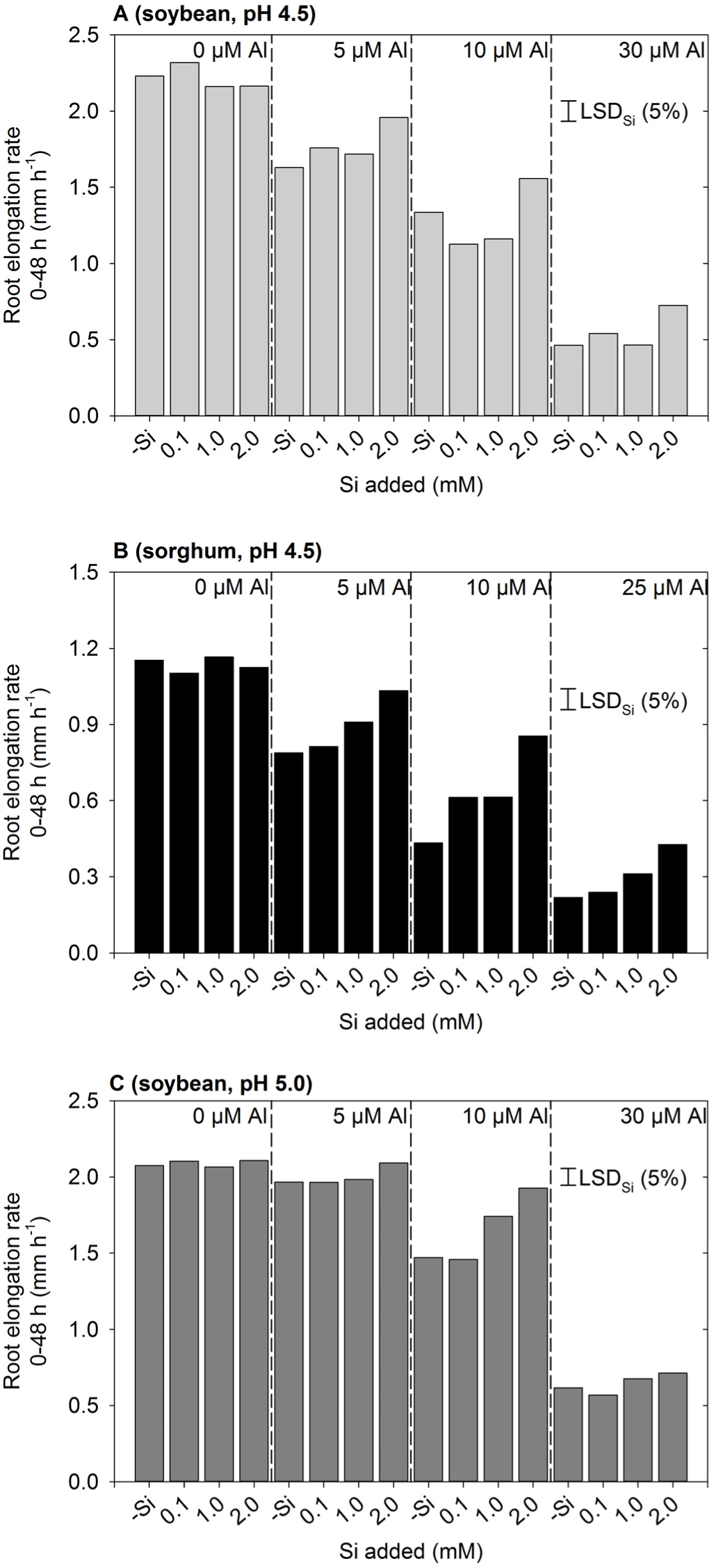
Effect of Al and Si on root elongation rate of (A) soybean in solutions at pH 4.5, (B) sorghum in solutions at pH 4.5, and (C) soybean in solutions at pH 5.0 (Experiment 1). The concentrations listed for Al and Si are nominal and were not measured. In all instances, the interaction between Al and Si was not significant, with the LSD values presented enabling the comparison between Si concentrations at any given Al concentration.

Experiment 2 aimed to provide more detailed information on the ability of Si to alleviate Al toxicity. Again, the addition of 2 mM Si was found to alleviate the toxic effects of Al, with RER being ca. 0.3 mm h^−1^ higher for soybean and ca. 0.2 mm h^−1^ higher for sorghum relative to values calculated from the regression in Si-free solutions (Figure 2). Using sorghum as an example, it was observed that RER in a solution with 25 μM Al and 2 mM Si (0.41 mm h^−1^) was comparable to that predicted for a solution with 13 μM Al and without added Si (as calculated from the regression, see dotted line in Figure 2B). Similar results were also observed for soybean (Figure 2A).

**Figure 2.**
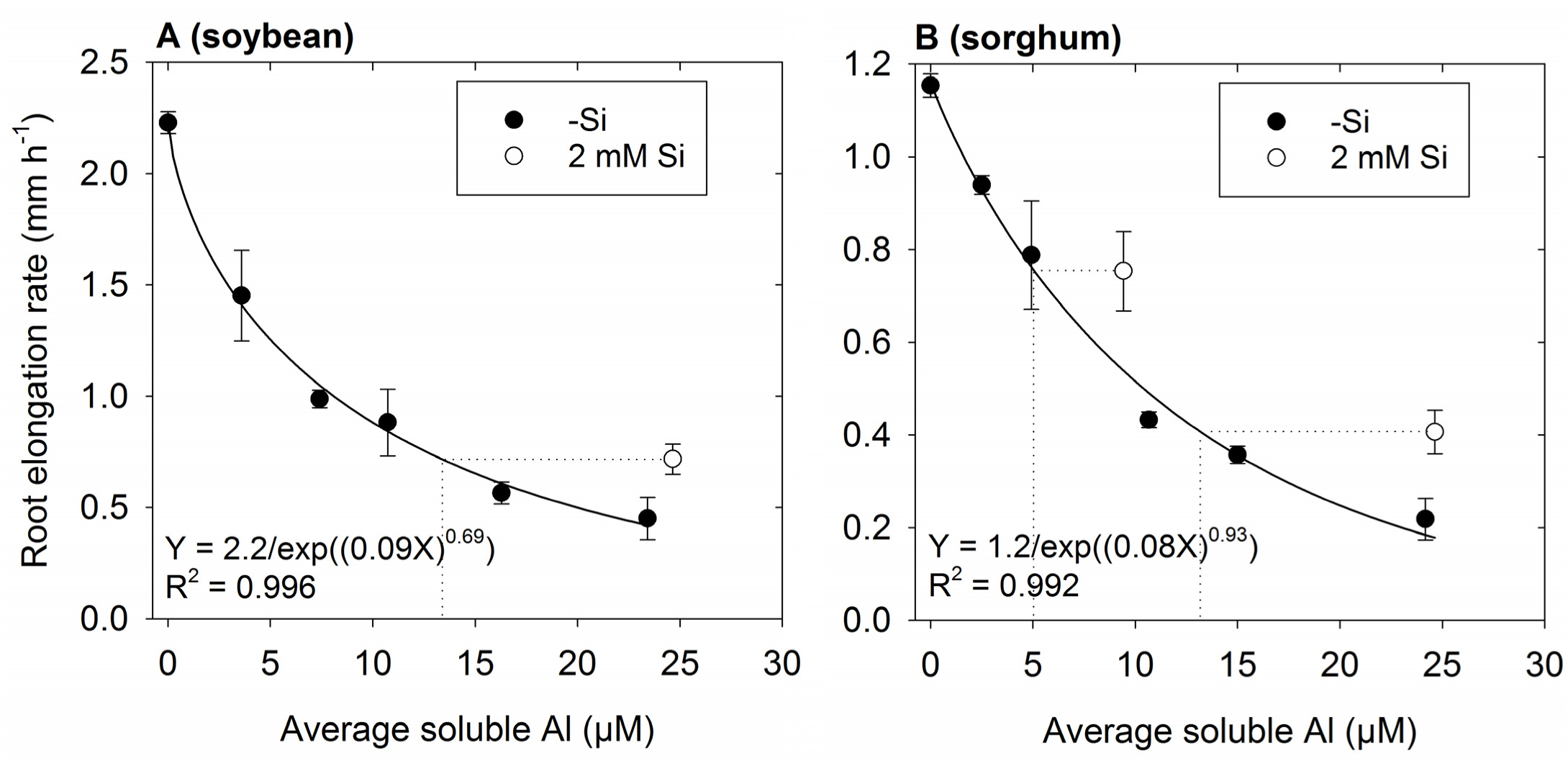
Effect of Al on root elongation rate (RER) of (A) soybean and (B) sorghum over 48 h exposure with either no added Si (-Si) or 2 mM Si at pH 4.5 (Experiment 2). Soluble Al concentrations are the average of values measured using the PCV method after 0, 24, and 48 h. The regression (Taylor et al., 1991) is fitted only to treatments with 0 μM Si. The dotted lines are shown for seedlings grown in solutions containing Si. These dotted lines show the predicted soluble Al concentration for a Si-free solution that corresponds to the same RER as observed in the solution containing Si.

To determine if this improved growth in Si-containing solutions resulted from a decrease in soluble Al due to the precipitation of aluminosilicate complexes within the bulk nutrient solution, inorganic monomeric Al concentrations were measured using the PCV method. Averaged across the 48 h experimental period, measured Al concentrations in Si-free solutions were an average of 12 % lower than the nominal concentrations, whilst measured Al concentrations in the three solutions with 2 mM Si were an average of 10 % lower than nominal concentrations (Figure 2). Thus, changes in RER upon the addition of Si were could not be attributed to changes in soluble Al concentrations. To further investigate potential changes in the Al chemistry of soluble Al in the bulk nutrient solution, we also modelled the potential formation of aqueous AlH_3_SiO_4_^2+^ using PhreeqcI (Table 1), with the magnitude of its formation predicted to be comparatively small. For example, for a solution containing 30 μM Al and 2 mM Si, aqueous AlH_3_SiO_4_^2+^ was predicted to have a concentration of 3.8 μM, with the predicted concentration of Al^3+^ concomitantly reducing from 23 μM (-Si) to 21 μM (2 mM Si). However, it should be noted that there is considerable uncertainty regarding the equilibrium constant for AlH_3_SiO_4_^2+^ (Browne and Driscoll, 1992; Pokrovski et al., 1996).

### 3.2 Effects of Si on concentrations of Al in root apical tissues

In the absence of Si, concentrations of Al in the apical root tissues (0-10 mm) increased with increasing Al in the nutrient solution (Figure 3). Although this was observed for both plant species, tissue Al concentrations were ca. three-fold higher in soybean than in sorghum. Importantly, for the Si-containing nutrient solutions, tissue Al concentrations were similar to those grown in the corresponding Si-free nutrient solutions (Figure 3).

**Figure 3.**
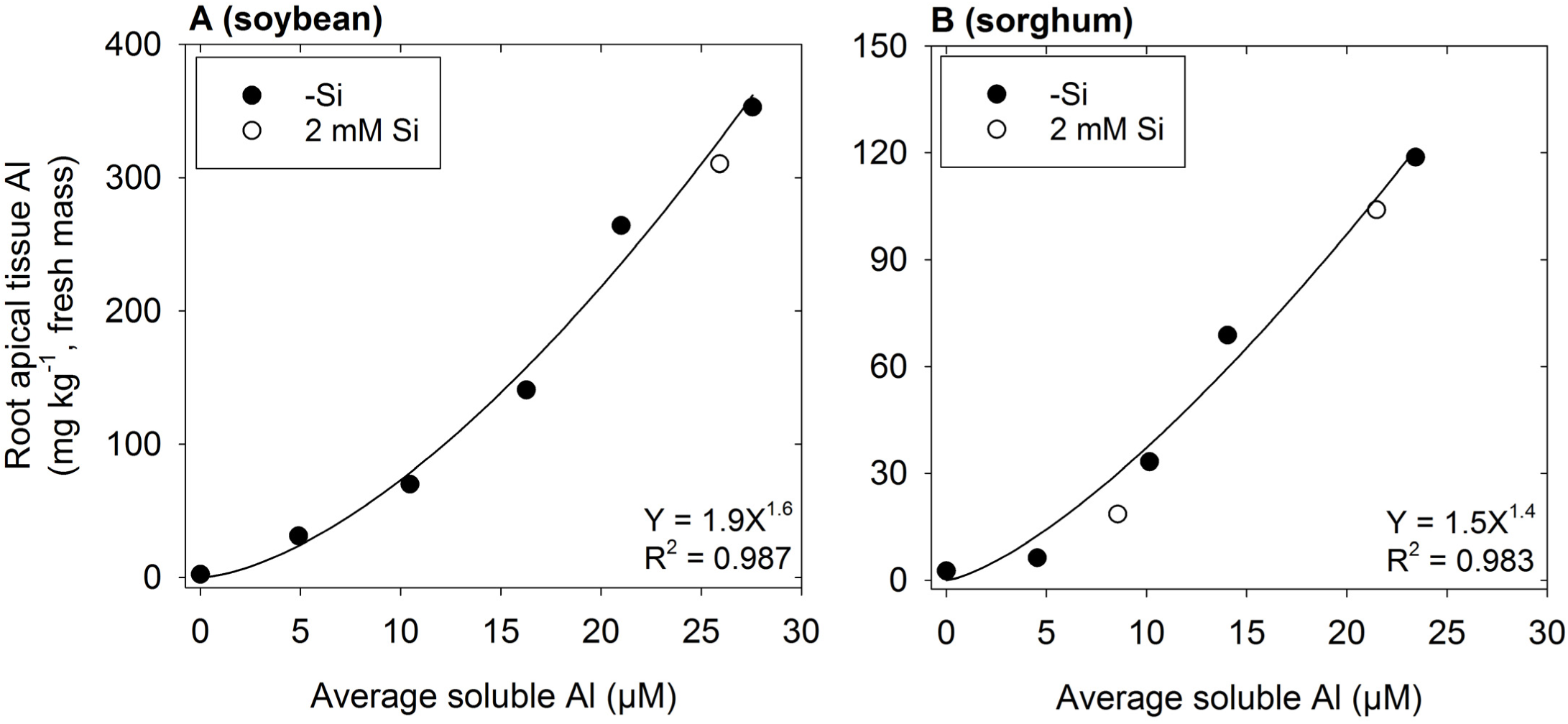
Effect of Al on the concentration of Al in the root apical tissues (0-10 mm) of (A) soybean and (B) sorghum. Soluble Al concentrations are the average of values measured using the PCV method after 0, 24, and 48 h. The regression is fitted only to treatments with no added Si (-Si).

### 3.3 *In situ* analyses of Al distribution in root apical tissues

To examine the cellular and subcellular distribution of Al and Si within the rhizodermis and outer cortex of the apical root tissues (5 mm from the apex) of sorghum, we utilized synchrotron-based LEXRF. In the absence of added Si, Al accumulated predominately in the cell walls (Figure 4A,B).In the presence of 2 mM Si, Al was again observed to accumulate in the cell walls, but some of this Al was co-located with high concentrations of Si (Figure 4C-F). These Al-Si complexes appeared to form either within the mucigel or in the outer apoplast at the root-solution interface, being associated both with the rhizodermis (Figure 4C,D) and with the border cells (Figure 4E,F).

**Figure 4.**
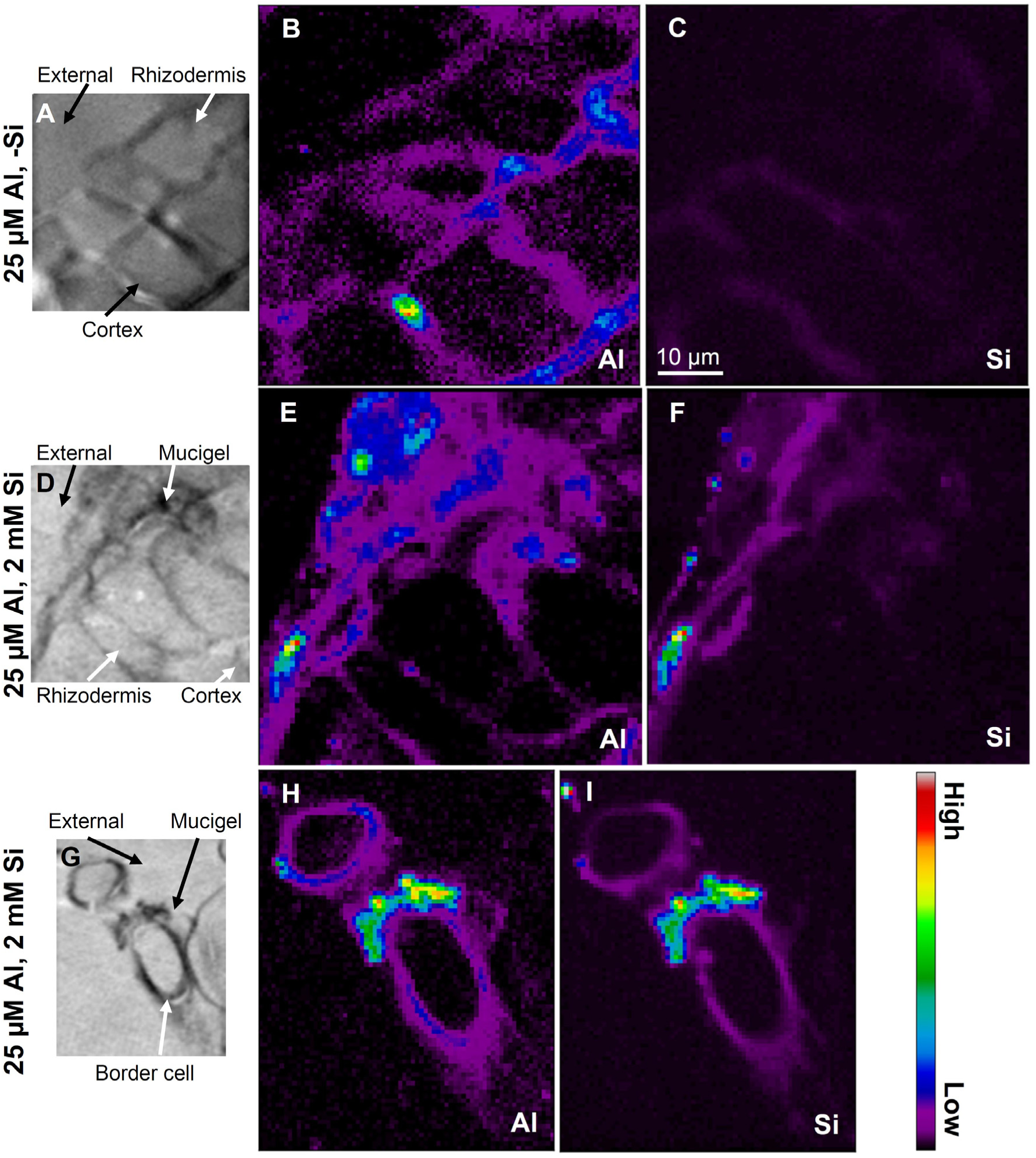
Distribution of Al and Si using LEXRF in 5-μm-thick transverse sections of sorghum roots (5 mm from the apex) exposed to solutions nominally containing either 25 μM Al (A-C) or 25 μM Al with 2 mM Si (D-I) for 48 h. The images in A, D, and G show the cellular structure, while images in B, E, and H show the distribution of Al, while images in C, F, and I show the distribution of Si. In (A-F), the exterior of the root is in the top-left corner, with the rhizodermis (and mucigel) plus 1-2 layers of cortical cells shown. Signal intensity is presented as a color scale, with brighter colors corresponding to higher concentrations. Images have been scaled so that they can be directly compared within each element [i.e. (B) can be compared with (E) and (H), and (C) can be compared with (F) and (I)], but not between elements. The scale-bar in (C) applies to all images.

Using the data from the LEXRF analyses, elemental correlation was examined to determine the relationship between Al and Si (if any). Unsurprisingly, in the absence of added Si, there was no relationship evident between Al and Si within the root tissues, with Si intensity being low for all pixels (Figure 5A). In contrast, in the presence of 2 mM Si, it was apparent that there were three different pixel ‘populations’, as shown in Figure 5B for example by the dashed ovals. For some pixels, although Al intensity was high, Si intensity remained low (blue dashed oval in Figure 5B). Conversely, some pixels had a low Al intensity but a high Si intensity (black dashed oval in Figure 5B, but also see Figure 5C). Finally, for some pixels, the intensity of both Al and Si was high, indicating the co-localization of these elements (red dashed oval in Figure 5B, but also see Figure 5C). Importantly, this increase with Si with increasing Al (red dashed oval in Figure 5B) was approximately linear.

**Figure 5.**
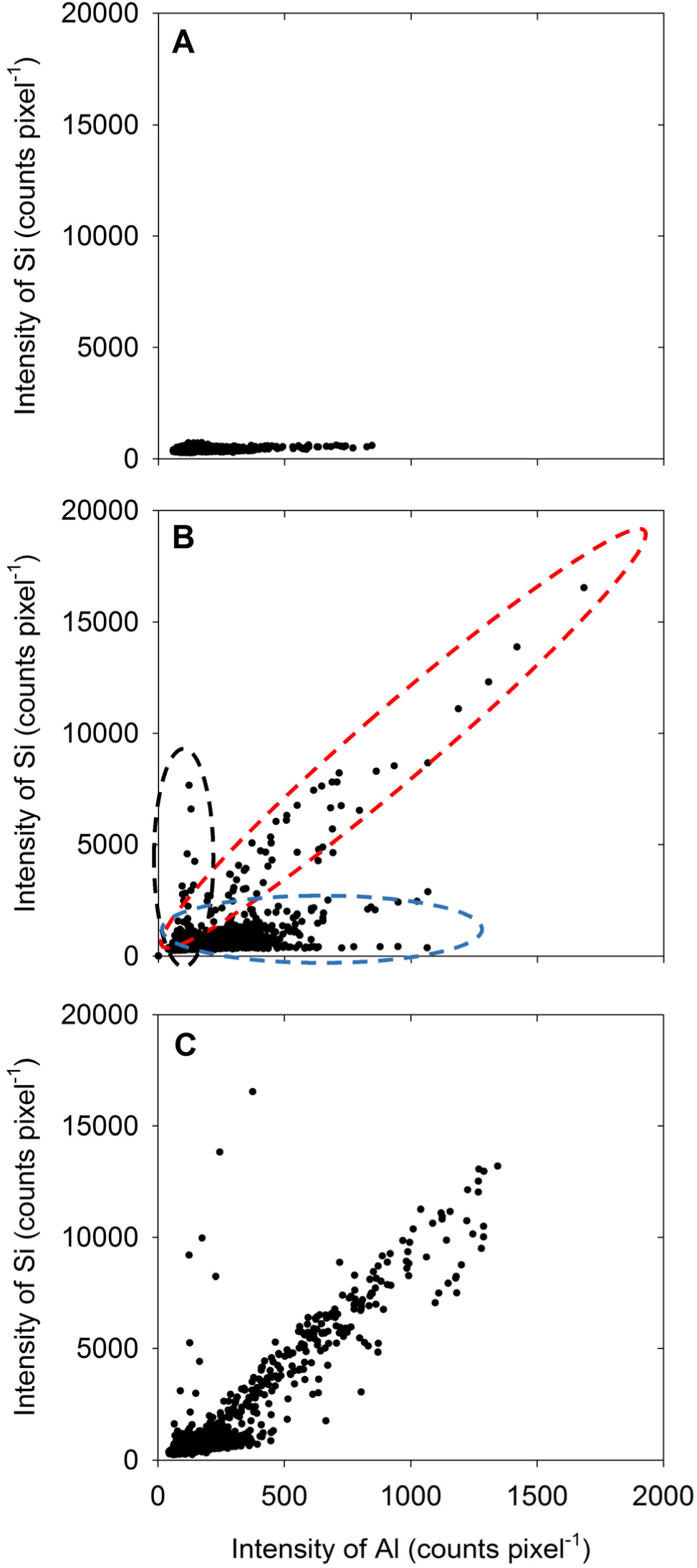
Element correlation plot showing the relationship between Al intensity and Si intensity for 5-μm-thick transverse sections of sorghum roots (5 mm from the apex) examined using LEXRF following exposure to solutions nominally containing either 25 μM Al (A) or 25 μM Al with 2 mM Si (B,C) for 48 h. Data were extracted from the images shown in Figure 4. Across the three samples (A-C), three broad types of populations were observed, as indicated by the three dashed ovals in (B).

## 4 Discussion

We found that the addition of Si partially alleviated the toxic effects of Al in both soybean and sorghum (Figure 1A,B and Figure 2). The magnitude of this Si-induced increase in RER in Al-toxic solutions was similar for both plant species, being up to ca. 0.3 mm h^−1^ (14 %) higher for soybean and ca. 0.2 mm h^−1^ (17 %) higher for sorghum relative to solutions without added Si. These findings are in agreement with that reported previously for soybean (Baylis et al., 1994) and sorghum (Galvez et al., 1987; Hodson and Sangster, 1993).

Although the addition of Si partially alleviated the reduction of root elongation in Al-toxic solutions (Figure 1A,B and Figure 2), this improvement in growth could not be attributed to a decrease in soluble Al in the bulk nutrient solution (Figure 2). Similarly, the improvement in growth could not be attributed to a decrease in the total concentration of Al in the apical (0-10 mm) root tissues (Figure 3). Using LEXRF, it was found that Al (in the absence of added Si) accumulated largely within the apoplast – a finding in accordance with previous observations in roots of soybean (Kopittke et al., 2015) and *Picea abies* (Hodson and Wilkins, 1991). For roots exposed to both Al and Si, the Al again accumulated in the apoplast, but much of the Al was co-located with Si both within the mucigel and within the apoplast (Figure 4C-F and Figure 5B,C). We consider it likely that these Al-Si complexes within the root tissues were aluminosilicates or hydroxyaluminosilicates, given their low solubility (Lindsay, 1979; Exley and Birchall, 1992), although further work is required in this regard. Thus, the data support the suggestion that Si alleviates Al toxicity through the *in planta* precipitation of Al-Si complexes. Given that Al rapidly reduces RER (within 5 min) by binding strongly to the cell wall and inhibiting its loosening (Jones et al., 2006; Horst et al., 2010; Kopittke et al., 2015), we propose that the formation of Al-Si compounds *in planta* reduces the toxic effects of Al by reducing its binding to the cell wall. This hypothesis is in accordance with the finding of Prabagar et al. (2011) who, examining suspension cultures of *Picea abies*, used morin (a fluorochrome) to examine the distribution of Al. Although it is known that morin does not bind to Al that is already bound strongly to other ligands (such as the cell wall) (Eticha et al., 2005), Prabagar et al. (2011) concluded that the addition of Si reduced the concentration of free Al within the cell wall in parallel with its amelioration of Al toxicity.

The magnitude of the improvement in RER upon addition of Si is in general agreement with previous observations, including in wheat (Cocker et al., 1998b), maize (Wang et al., 2004), and sorghum (Galvez et al., 1987; Hodson and Sangster, 1993) [also see Table 1 of Hodson and Evans (1995)]. In addition, our finding that the Si-induced improvement is associated with the formation of Al-Si compounds in the mucigel or within the outer apoplast at the root-solution interface are in general agreement with those of Hodson and Sangster (1993). These authors used electron microscopy to examine roots of sorghum exposed to 100 μM Al and 2800 μM Si for 8 d and found Al-Si compounds in the outer tangential wall (i.e. at the root-solution interface). The solution Al concentration of 100 μM used by Hodson and Sangster (1993) however, is high relative to that commonly found in solutions of acid soils (Kopittke and Blamey, 2016) and is also high relative to that required to reduce RER in sorghum (Figure 2). It is possible that Hodson and Sangster (1993) used this high Al concentration due to the comparatively poor sensitivity of the energy-dispersive X-ray spectroscopy that was used to analyze the plant tissues. Our findings are also in agreement with those of Wang et al. (2004), with these authors using a fractionation procedure and reporting that the addition of Si resulted in greater accumulation of Al in the root cell walls, possibly as aluminosilicates. In contrast, however, the findings of the present study do not seem to be in agreement with Corrales et al. (1997) who, studying maize, found that the addition of Si decreased Al concentrations in root tissues, with these authors proposing that this decrease in Al concentration in the root tissue was possibly associated with decreased binding to cell wall through esterification of polyuronides within cell walls (Corrales et al., 1997). Similarly, it was suggested by Kidd et al. (2001) that an enhanced exudation of phenolic compounds may be responsible for Si-induced Al resistance in maize.

It is useful to consider why these Al-Si compounds formed within the mucigel or apoplast but not in the bulk nutrient solution (which would have resulted in a decrease in the inorganic monomeric Al concentration). Firstly, it is known that the mucigel is a highly negatively charged polysaccharide (having the highest negative charge of any portion of the root) as is the cell wall (Oades, 1978; Haynes, 1980). Therefore, the observed binding of Al to the mucigel and cell wall (Figure 4) is expected and in accordance with previous observations (Hodson and Wilkins, 1991; Kopittke et al., 2015). However, the formation of Al-Si complexes within the mucigel or apoplast presumably occurred due to supersaturation resulting from the increased Al and Si concentrations within these root tissues (Figure 4 and Figure 5) together with an increased pH. For example, it is has been reported that the pH of the root apoplast is generally 5-6 (Staal et al., 2011), compared to the bulk solution pH value of 4.5 in the present study. Indeed, it is known that the solubility of aluminosilicates and hydroxyaluminosilicates decrease as pH increases towards neutral (Lindsay, 1979; Exley and Birchall, 1992), and it is well known that hydroxyaluminosilicates form in neutral and slightly alkaline solutions.

In conclusion, in accordance with previous studies, we have found that Si can potentially increase RER in Al-toxic solutions, with the magnitude of the effect increasing with the concentration of Si. This improvement in RER could not be attributed to a change in Al-chemistry of the bulk nutrient solution, with inorganic monomeric Al concentrations remaining constant despite the addition of Si at concentrations of up to 2 mM. Similarly, the improvement in RER could not be attributed to a decrease in the concentration of Al within the apical root tissues. We used LEXRF to examine the distribution of Al and Si within the rhizodermis and outer cortex in tissues 5 mm from the root apex. It was found that Al accumulated in the cell wall and mucigel, but that in roots also exposed to Si, high concentrations of Al were observed to be co-located with Si in the mucigel and outer apoplast at the root-solution interface. Thus, the present study has shown that Si can reduce the toxicity of Al *in planta* through formation of Al-Si complexes, with this presumably decreasing the strong binding of Al to the cell wall where it exerts its toxic effects. Given that Si is the second most abundant element in the earth’s crust, the findings of the present study have important implications for understanding the toxic effects of the elevated levels of Al that occur in the estimated 30-40 % of all arable soils worldwide are acid.

## 6 Author Contributions

P.M.K. conceived the research program. P.K.K., K.G., and B.A.McK. conducted the plant growth experiments; P.M.K., A.G., G.K., and B.A.McK. conducted LEXRF analyses at Elettra (Italy); P.M.K. wrote the first draft of the article to which all other authors contributed.

## 8

### Acknowledgments

The assistance of Professor Chris Exley (Keele University, UK) is appreciated.

